# An *in vivo* massively parallel platform for deciphering tissue-specific regulatory function

**DOI:** 10.1101/2022.11.23.517755

**Authors:** Ashley R. Brown, Grant A. Fox, Irene M. Kaplow, Alyssa J. Lawler, BaDoi N. Phan, Morgan E. Wirthlin, Easwaran Ramamurthy, Gemma E. May, Ziheng Chen, Qiao Su, C. Joel McManus, Andreas R. Pfenning

## Abstract

Genetic studies are rapidly identifying non-protein-coding human disease-associated loci. Understanding the regulatory mechanisms underlying these loci remains a challenge because the causal variants and the tissues in which they act are often unclear. Massively parallel reporter assays (MPRAs) have the potential to link differences in genome sequence, including genetic variants, to tissue-specific regulatory function. Although MPRA and similar technologies have been widely adopted in cell culture, there have been several barriers to widespread use in animals. We overcome these challenges with a new whole-animal MPRA (WhAMPRA), where systemic intravenous AAV effectively transduces the plasmid MPRA library to mouse tissues. Our WhAMPRA approach revealed models of tissue-specific regulation that generally match machine learning model predictions. In addition, we measured the regulatory effects of disrupting MEF2C transcription factor binding sites and impacts of late onset Alzheimer’s disease-associated genetic variations. Overall, our WhAMPRA technology simultaneously determines the transcriptional functions of hundreds of enhancers *in vivo* across multiple tissues.

## Main Text

Transcriptional regulation, a process in which non-coding enhancer sequences play a major role, is a key component of specifying both cell type identity and phenotypic diversity ^1–4^. In neural tissue, gene regulatory processes are essential for organizing the range of highly interconnected and regionally specialized cell types that must synchronize their activity to produce behavior ^5^. Transcription is largely regulated by enhancers, distal non-coding sequences that are highly tissue-specific relative to proximal promoters ^6^. Recent progress in experimental technology has allowed direct profiling of open chromatin, a component of the “epigenomic” (i.e. gene regulatory) landscape, both at the level of tissues as well as individual cell types ^7–11^. In parallel, advances in computational methods have enabled the development of sophisticated algorithms, trained on these open chromatin datasets, for predicting the regulatory activity of enhancers from genome sequence alone ^12–14^.

High-throughput reporter assays ^15^ can be used to experimentally validate and refine the predictions of machine learning models of gene regulatory activity by measuring the impact of genetic differences across species on regulatory element activity *in vivo*. Chief among these methods is the massively parallel reporter assay (MPRA) ^16–20^, which involves creating a library of thousands of distinct plasmids, each of which contains a custom-synthesized candidate enhancer element that controls the expression of a unique barcode in conjunction with a minimal promoter (**Fig. 1A**). MPRA libraries can be constructed using enhancer capture as well as other methods ^21–23^. In these studies, the expression of these unique plasmid barcodes as RNA, which can be measured in parallel by complementary DNA (cDNA) amplicon sequencing, reflects the transcriptional activity of the corresponding enhancer in the particular cells into which the library has been introduced.

**Figure 1.**
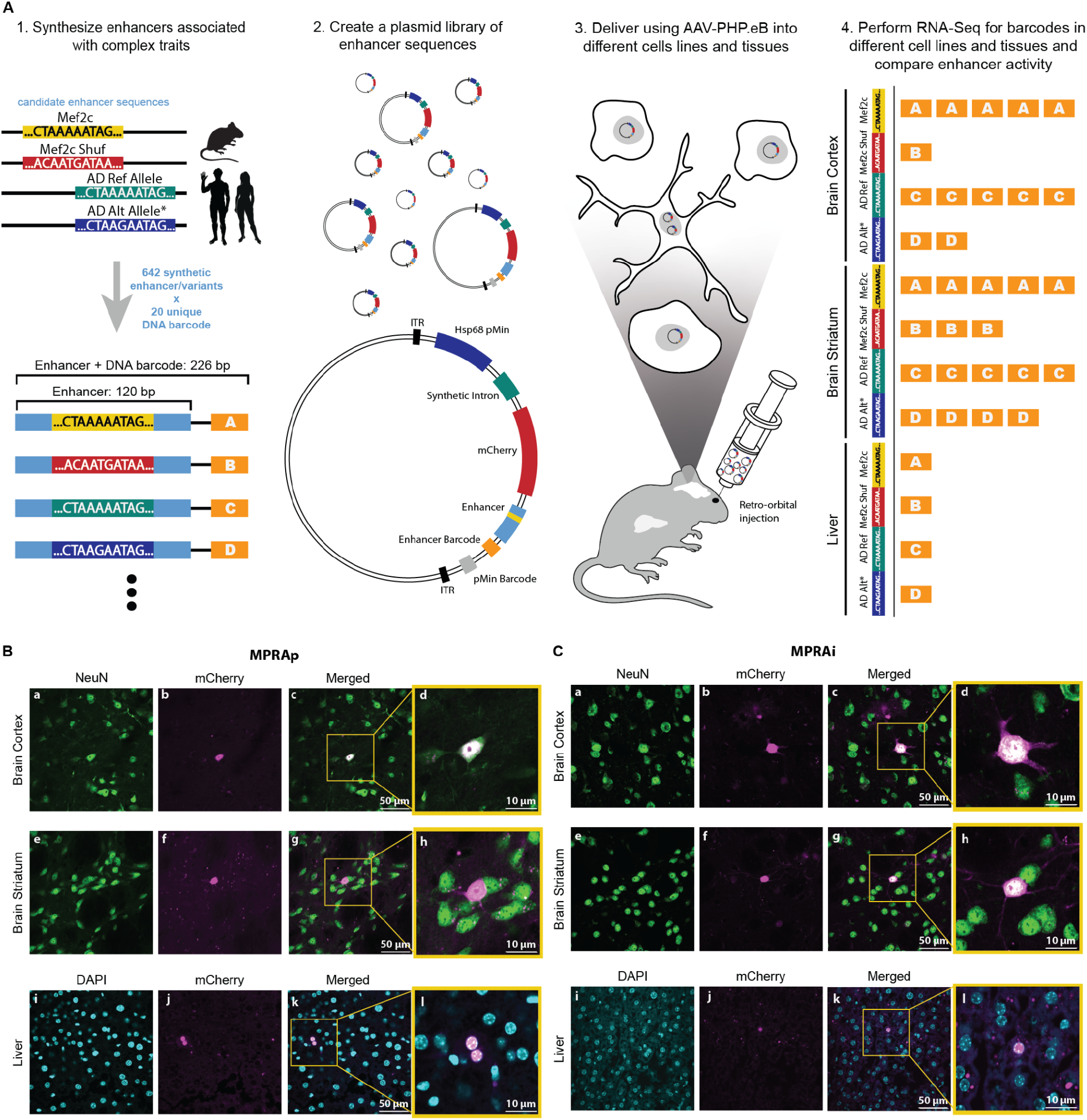
WhAMPRA tests effects of transcription factor binding and single nucleotide variations on transcriptional regulation. (A) 1. The library is designed to study complex traits consisting of 642 enhancers and variants, each with 20 unique barcodes (i.e. MEF2C motifs and shuffled versions of these motifs, as well as reference and alternative alleles for AD-associated SNPs). 2. The oligos are synthesized and cloned into plasmids. 3. The plasmid library is packaged into the PHP.eB AAV serotype and delivered into a mouse via retro-orbital injection. 4. The activity of candidate enhancers in multiple brain regions and tissues is measured using RNA levels of the bar-codes. (B) mCherry expression from MPRAp (cross-tissue positive controls). Shown is mCherry (magenta) compared to NeuN expression (green) in the brain cortex (panels a-d) and brain striatum (panels e-h) from a C57Bl/6J mouse. mCherry (magenta) is also compared to DAPI (blue) expression in the liver (panels i-l) from a C57Bl/6J mouse. (C) mCherry expression from MPRAi (MPRA library of 642 enhancers/variants). Shown is mCherry (magenta) compared to NeuN expression (green) in the brain cortex (panels a-d) and brain striatum (panels e-h) from C57Bl/6J mouse. mCherry (magenta) is also compared to DAPI (blue) expression in the liver (panel i-l) from a C57Bl/6J mouse.

The ability to effectively deliver plasmid libraries of hundreds-to-thousands of candidate enhancers to cultured cells has allowed for detailed, quantitative descriptions of how subtle differences in genome sequence relate to differences in cell type-specific gene regulation. Studies have used reporter assay techniques to measure the effects of single nucleotide polymorphisms (SNPs) identified from expression quantitative trait loci studies (eQTLs) ^19,20^, SNPs from genome-wide association studies (GWAS) ^24–26^, and human lineage-specific mutations ^27,28^. High-throughput reporter assays have also been adapted to study gene regulation in cultured neurons ^18,29^. However, cultured neurons fail to capture the full complexity of highly connected and regionally specialized *in vivo* neural tissue ^30,31^.

MPRAs have also been adapted to study gene regulation in the brain *in vivo*, but the current assays lack the sensitivity to detect subtle differences in activity from disrupting individual transcription factor binding sites or SNPs. Both electroporation and adeno-associated viruses (AAVs) have been used to deliver MPRA libraries to the mouse brain ^23,32–34^. Although these studies have the sensitivity to measure the tissue or even cell type-specificity of enhancers ^35^, limitations in the number of cells receiving the libraries has made it impossible to detect the impact of genetic variants on brain gene regulation ^33,36^.

To this end, we developed a whole-animal massively parallel reporter assay (WhAMPRA), which combines a custom, highly modular plasmid with the AAV-PHP.eB virus ^37^ to deliver the plasmid library containing our reporter assay to a broad range of tissues within a single animal with high reproducibility (**Fig. 1**). With a brief, minimally invasive intravenous injection rather than direct injections, this serotype enters the brain by crossing the blood-brain barrier, increasing the breadth and the consistency of expression across the tissue. This also enables direct comparisons within a single mouse of enhancer activity between different brain regions and the brain versus other tissues. We demonstrate that WhAMPRA can measure the effects of synthetic disruptions of candidate transcription factor binding sites and naturally occurring human variants on tissue-specific enhancer activity.

## Results

### WhAMPRA libraries are successfully delivered to tissues across the mouse

We designed the WhAMPRA plasmid (pAAV-MPRAe) and delivery system to maximize transduction and reproducibility. We designed the vector with an Hsp68 minimal promoter, which can stabilize the enhancer-promoter interaction, assisting the initiation of transcription. We also included an mCherry reporter. We cloned the synthesized candidate enhancer sequences and barcodes downstream of both the minimal promoter and mCherry. Cloning sites within the plasmid backbone allow modular changes to both promoter and enhancer sequences (**Supplementary Fig. 1**).

We first tested the transduction and expression of our reporter system by cloning 3 cross-tissue positive control sequences each with one barcode (**Supplementary Table 1**) into the WhAMPRA plasmid after the mCherry coding sequence to make a small MPRA library (MPRAp) followed by transduction into wildtype adult mouse tissue. We confirmed successful transduction and transcription tissues of interest, including neurons in multiple brain regions, by imaging for mCherry (**Fig. 1B**).

We then designed a library of 642 enhancers each paired with 20 unique barcodes (MPRAi) to test the ability of WhAMPRA to relate differences in genome sequence to regulatory differences at multiple levels (**Fig. 1A and Supplementary Table 2**). First, we selected a set of expected positive and negative controls from brain, liver, and immune cells ^17,18^ based on their measured regulatory activity in previous MPRA experiments. To improve the overall signal from mouse brain, we also added 144 candidate enhancers based on mouse cortex H3K27ac chromatin immunoprecipitation sequencing (ChIP-seq) regions and their orthologs across species (see methods). To test the ability of the assay to detect the impact of disrupting transcription factor binding sites, we also synthesized versions of 28 sequences that are known to bind the transcription factor MEF2C, which has been implicated in transcriptional regulation in multiple brain regions ^38,39^ as well as Alzheimer’s disease (AD) predisposition ^40^. Finally, we synthesized different versions of a set of 27 candidate enhancers with both the risk and the non-risk alleles of candidate regulatory AD-associated variants from GWAS ^41^.

We cloned the candidate enhancer sequences into the plasmid backbone and then delivered the libraries into the brains of mice using retro-orbital injection of AAV-PHP.eB ^37^ as described in Lawler *et al*. ^42^ (**Fig. 1A**). We collected tissue and sectioned from the liver, the frontal cortex, and the striatum. Imaging nuclear mCherry relative to NeuN (brain) or DAPI (liver) revealed widespread systemic transduction (**Fig. 1B**). Given that not every candidate enhancer is active in every tissue and enhancer activity varies, exact transduction rates are better measured by sequencing of plasmid DNA counts rather than by imaging.

We next sought to catalog the transduction within our library by using DNA sequencing to measure the levels of unique barcodes in cells across tissues. We injected libraries into 8 mice and collected samples from the liver, the primary motor cortex (M1), a larger piece of tissue from the rest of the frontal cortex (referred to as cortex), the hippocampus, and the striatum. We also collected a few samples of tissue from the heart, kidneys, testes, and ovaries. To compare *in vivo* and cell culture technologies, we also transfected our library in the microglia-like HMC3 cell line. The use of HMC3 also allowed us to study the potential function of MEF2 binding sites and AD-associate genetic variants, which have both been linked to microglia ^43,44^. Measurements of the plasmid DNA from each sample were highly correlated (Spearman Rho ranging from 0.737 to 0.991, median 0.951; **Supplementary Fig. 2A and Supplementary Table 3**). Most samples had a high proportion of barcodes detected at the DNA level (**Supplementary Figure 2B**). These results confirm widespread transduction across mouse tissues.

### WhAMPRA measures the tissue-specificity of candidate enhancers

The estimated enhancer activity levels suggest that WhAMPRA can reliably measure regulatory activity across different tissues. We used RNA-Seq to measure barcode RNA expression at the RNA levels in HMC3 cells, the liver, the M1, cortex, the hippocampus, and the striatum (**Fig. 1A, right and Supplementary Table 4, 5**). The RNA barcodes showed less reproducibility across samples than the DNA barcodes, likely due to the tissue-specificity of gene regulation (Spearman Rho ranging from 0.366 to 0.995, median 0.629; **Supplementary Fig. 2C and Supplementary Table 3**). Most samples had a high proportion of barcodes detected at the RNA level, but samples with the highest levels came from brain, HMC3, and liver, the tissues for which the enhancers were designed (**Supplementary Figure 2D**).

We used the barcode RNA counts relative to the barcode DNA counts to estimate enhancer activity with MPRAnalyze ^45^. We found that the candidate enhancers, which include positive controls, had a strong tendency to be expressed relative to the negative control sequences (**Fig. 2A**). The strongest enrichments for active enhancers came from the brain, where most of the candidate enhancers were expected to be active. Quality control metrics were consistent between the *in vivo* and *in vitro* version of the experiment (**Supplementary Table 3**). For example, RNA:DNA ratios from the HMC3 cells (**Fig. 2B**) showed a similar spread to the RNA:DNA ratios from brain tissue like cortex (**Fig. 2C**).

**Figure 2.**
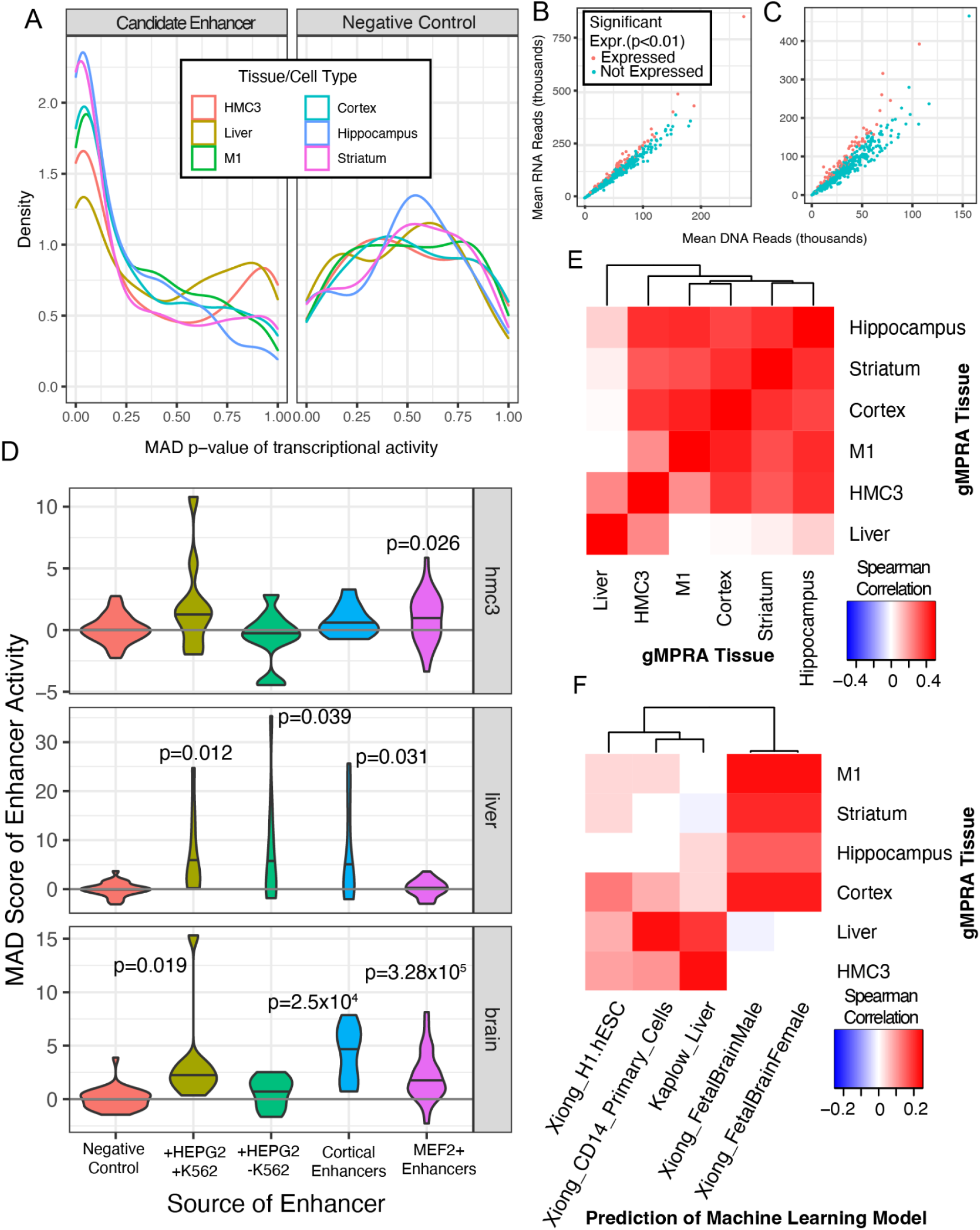
WhAMPRA captures tissue-specific signatures of gene regulation *in vivo*. (A) The frequency of p-values is displayed using a density plot across the candidate enhancers and positive controls (left) relative to the negative controls (right). The ratio of DNA reads to RNA reads, which roughly corresponds to transcriptional activity, is plotted for HMC3 cultured cells (B) and for cortical tissue (C). The mean across all samples for that tissue is used. (D) The MAD score is displayed as a violin plot for the likely positive and negative control enhancers gleaned from other MPRA experiments. (E) Spearman’s Rho is calculated across the estimated transcription rate, alpha, of all enhancers for each pairwise tissue comparison. (F) Spearman’s Rho is calculated between the estimated transcription rate, alpha, of all enhancers and the prediction of open chromatin levels calculated by the convolutional neural network models.

The activity of the control enhancers provides evidence that WhAMPRA is able to identify tissue-specific patterns. Relative to the negative control candidate enhancers, the set of enhancers active in both HEPG2 (liver-like) and K562 (immune) cells ^17^ showed a nominal trend toward expression in HMC3 cells (one-sided t-test p=0.078), the liver (one-sided t-test p=0.012), and the brain tissues (one-sided t-test p=0.019; **Fig. 2D**). In contrast, the HEPG2-specific enhancers tended to be transcribed in only the liver (one-sided t-test p=0.039). The cortical control enhancers showed the highest regulatory activity in the brain (one-sided t-test p=0.00025; **Fig. 2D**). Overall, the expression patterns of the positive and negative control candidate enhancers matched expectations. Thus, our experimental design identifies tissue-specific differences in enhancer activity in living mice.

The patterns of expression across all enhancers also provide evidence that the WhAMPRA can capture tissue-specific gene regulation. There was strong correlation between enhancer activity across different brain tissues (Spearman Rho 0.348 to 0.433) but little correlation between brain and liver enhancer activity (Spearman Rho 0.0018 to 0.0971; **Fig. 2E**). The microglia-like cell line, HMC3, showed similarities to both brain and liver tissue (Spearman Rho 0.225 to 0.407) (**Fig. 2E**). To ensure that the enhancer activity we were measuring was related to the tissue-specific regulatory code, we compared the measured activity across all enhancers to machine learning model predictions of open chromatin ^12,14^, which are correlated with enhancer activity ^6^ (**Supplementary Table 6**). The predictions of machine learning models trained on brain tissue open chromatin had significant correlations with the WhAMPRA-measured enhancer activity in brain tissue (Spearman Rho 0.121 to 0.183; p from 9.39×10^−3^ to 7.90×10^−5^) but not liver tissue (Spearman Rho =-0.0117, -0.00370; **Fig. 2F**). Reciprocally, the predictions from a machine learning model trained on liver open chromatin were correlated with enhancer activity in liver (Spearman Rho=0.158; P=6.68×10^−4^) but not brain (Spearman Rho from -0.0112 to 0.0341; **Fig. 2F**). These results show that WhAMPRA identifies tissue-specific signatures of transcriptional enhancers in live mice.

### WhAMPRA detects enhancer disruptions from transcription factor binding sites and SNPs

To determine whether WhAMPRA could detect how disruption in transcription factor binding site motifs disrupts enhancer activity *in vivo*, we designed a set of 28 candidate enhancer sequences based on binding the MEF2C transcription factor in mouse cortex (see methods).

We also created versions of each enhancer where the transcription factor binding site itself was shuffled, along with an additional version where the motif together with the surrounding 5 nucleotides were shuffled (**Fig. 3A**). The non-disrupted MEF2 motif-containing enhancers showed the strongest activity in brain tissue but also showed some activity in HMC3 cells, consistent with MEF2C’s function in both brain and microglia ^38,43,46^ and its lack of expression in the liver (**Fig. 2D**) ^47,48^. Consistent with that observation, we found that disrupting the MEF2C transcription factor binding sites had no significant effect in liver tissue (p>0.1; **Fig. 3B, top**) but significantly decreased enhancer activity in HMC3 cells and in cortical tissue (one-tailed t-test p-value = 0.002; **Fig. 3B, middle and bottom**). Notably, the extent to which enhancer activity was disrupted by shuffling the MEF2C binding site was highly correlated with the original baseline expression of the enhancer (**Fig. 3C; Supplementary Table 7**; Rho=0.88; p=9.3×10^−7^). This suggests that the cases where disrupting the MEF2C transcription factor binding site has no effect are cases where the enhancer itself is not active in the assay rather than cases where the MEF2C binding site is not important for enhancer activity.

**Figure 3.**
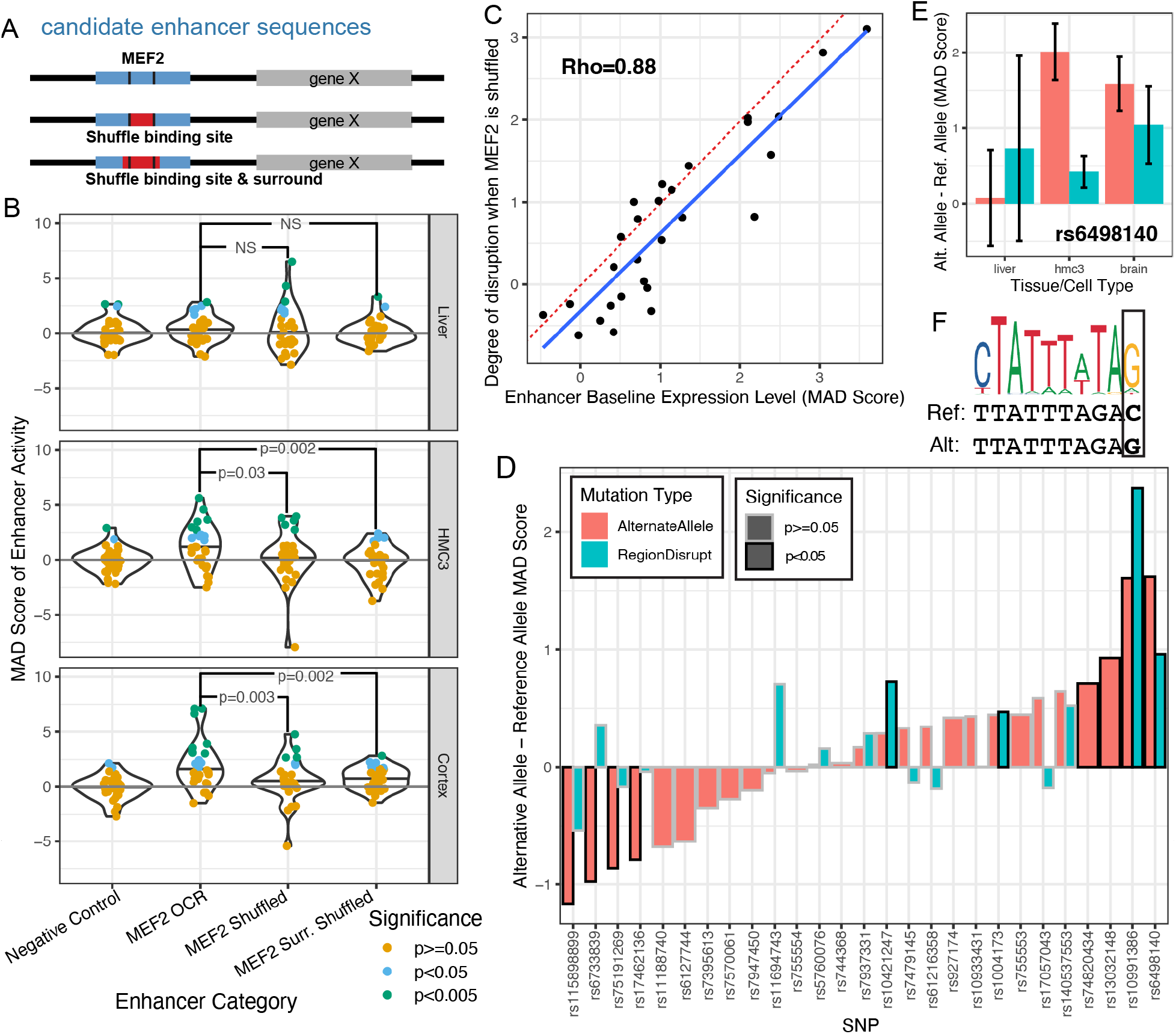
WhAMPRA detects enhancer differences due to MEF2C binding site disruption and candidate Alzheimer’s disease SNPs. (A) The experimental design of how MEF2C was systematically disrupted at candidate enhancers with binding sites. (B) The MAD score of enhancer activity is compared between negative control enhancers and different versions of candidate MEF2C enhancers. Each enhancer is colored based on its nominal significance of transcription relative to the population of negative controls (MAD p-value). (C) The MAD score of baseline enhancer expression is compared to the difference between the baseline enhancer expression and the average expression across the two instances of MEF2C shuffling. (D) The MAD score of the enhancer activity from the reference allele is compared to the alternate allele (red) and the sequence with a shuffled local transcription factor binding site (blue). (E) The MAD score of the alternative allele for SNP rs6498140 is compared to the MAD score of the reference allele for the candidate AD-associated enhancer. (F) The motif logo for a discovered MEF2 transcription factor binding site is visualized above the reference and alternative allele for rs6498140.

Next, we used WhAMPRA to measure the impact that 27 Alzheimer’s Disease GWAS-derived SNPs have on regulatory activity. We synthesized both the risk and non-risk allele of non-coding SNPs implicated in an AD genome-wide association study ^41^. For the subset of those enhancers where the SNP disrupted an important transcription factor binding site ^49^, we also included a version of the enhancer where the entire transcription factor binding site was shuffled. There were eight enhancers where the two alleles showed differential activity across the HMC3 and brain samples, two of which (rs6498140, rs10991386) were confirmed by the motif disruption (**Fig. 3D**; **Supplementary Table 8**).

The strongest evidence for the impact of a candidate AD-associated SNP on enhancer activity is for rs6498140, which is proximal to the gene *CLEC16A*. The alternate allele has the highest regulatory activity in both brain tissue and in HMC3 (**Fig. 3E**). The alternate allele creates a MEF2 transcription factor binding site motif in the enhancer (**Fig. 3F**), which is consistent with MEF2 being an active transcription factor in both brain and microglia ^43^. Notably, the SNP rs6498140 displays marginal GTEx eQTL associations in several tissues including the frontal cortex, where the alternate allele correlates with higher expression of *CLEC16A* ^50^Thus, our WhAMPRA both validates and expands the understanding of genetic variation and gene regulation in AD *in vivo* at relevant tissues.

## Discussion

While MPRA technology has been instrumental in linking genome sequence to regulatory function, thus far it has been used primarily in cell culture. Here, we developed WhAMPRA, a massively parallel reporter assay with the ability to measure regulatory activity across tissues *in vivo*. In concert with other experimental design choices, the delivery of the MPRA library into cells using systemic AAV transduction helps us achieve the necessary sensitivity to detect the tissue-specific regulatory effects of mutating transcription factor binding sites and individual nucleotides. Systemic delivery reduces local inflammation caused by intraparenchymal injection and thereby enables the study of disease processes with inflammatory components ^51,52^. Furthermore, systemic delivery increases the throughput of delivering MPRA to multiple tissues of interest in the same experimental animal with one brief, minimally invasive procedure.

A comparison of regulatory activity in brain and liver demonstrates that WhAMPRA can detect highly tissue-specific regulatory activity. We then use candidate enhancers known to bind the MEF2C transcription factor, which is active in the brain in microglia, to show that the assay can detect the impact of disrupting MEF2C transcription factor binding sites on regulatory activity. We also use a set of candidate AD-associated mutations to show that the assay is sensitive enough to detect the impact of disrupting individual SNPs on enhancer activity.

Although we are able to detect tissue- and allele-specific effects, there are still limitations in WhAMPRA’s *in vivo* technology. In contrast to the hundreds of enhancers we profile, current cell culture reporter assays can provide quantitative, cell line-specific information across thousands of enhancers ^17,27,53^. Efficient transduction of tissue is likely the current limitation in the technology. Furthermore, any AAV tropism present in PHP.eb or another systemic delivery system will also be reflected in the measured enhancer activity. For example, gene regulatory programs active in brain microglia are not likely to be captured in our WhAMPRA experiment due to the bias that PHP.eb has for neurons and other glial cells ^37^.

As AAV technology improves, we expect WhAMPRA to become more flexible. We designed our library to have active enhancers in brain, liver, and immune cells, but we detect transduction in heart, kidney, and other other tissues. New AAV variants are being designed that would allow WhAMPRA libraries to be targeted to specific cell subtypes ^54,55^ and even non-human primate tissue ^56^. As transduction efficiency improves, WhAMPRA could be paired with a methodology to isolate individual cell types ^9,42^ or even single-cell profiling ^32^.

GWAS and whole-genome sequencing studies are identifying an increasing number of candidate regulatory variants underlying the predisposition to complex traits. Fine-mapping and functional characterization of those variants is an important step in connecting genetic predisposition to disease pathophysiology. While *in vitro* high-throughput reporter assays have provided an avenue for high-throughput functional characterization in human cells, there may be genetic variants that act in a cell type or cellular environment not able to be captured *in vitro*. Complementary to *in vitro* versions of MPRA, WhAMPRA enables high throughput functional characterization across tissues of a behaving organism. This enables tissue-specific regulatory effects to be measured in animal models of disease, including non-traditional model organisms.

## Acknowledgements

One or more of the authors of this paper self-identifies as a member of the LGBTQ+ community. We thank Drs. Viviana Gradinaru and Benjamin Deverman for early use of their systemic brain-transducing viral capsid. We thank Dr. Jesse Gray and the members of his lab for their discussion of early versions of WhAMPRA. We thank Dr. Oliver Schlüter and Corinne Schneider from the University of Pittsburgh for their training and discussions about AAV production. We thank Tae Yoon Park and other current and former members of the Pfenning lab as well as the CMU Neuroscience community, especially Drs. Alison Barth, Aryn Gittis, and Sandra Khulman, for their discussions and feedback throughout this project. MEW was funded by the Carnegie Mellon BrainHub Postdoctoral Fellowship. IMK was funded by the Carnegie Mellon Computational Biology Department Lane Fellowship. BNP was funded by the NIH National Institutes on Drug Abuse (F30DA053020). AJL was funded by the NSF (DGE1745016). ARP was funded by the NIH National Institutes on Drug Abuse (DP1DA046585) and the CURE Alzheimer’s Fund.

