## Supplementary Material for "An *in vivo* massively parallel platform for deciphering tissue-specific regulatory function"

**Affiliations:** Carnegie Mellon University Departments of [1]Computational Biology and [2]Biology and [3]Neuroscience Institute; [4]Medical Scientist Training Program, University of Pittsburgh School of Medicine; [5]Current affiliation: Stanley Center for Psychiatric Research, Broad Institute; [6]Current affiliation: Allen Institute for Brain Science; [7]Current affiliation: College of Medicine, University of Kentucky

### Supplemental Online Methods:

#### Array design:

Cross-Tissue (CT) Positive Controls

The goal of global MPRA is to produce robust delivery of plasmid expression vectors to a tissue of interest using AAV. Since transduction and transcription of episomal plasmid DNA are both properties of the virus serotype and regulatory element, we diversified our potential to identify tissues where WhAMPRA would work by using 3 viral promoters and enhancers. We selected the 72bp SV40 enhancer element; the 245bp SV40 promoter, which includes 1 copy of the 72bp enhancer [57]; and the 305bp CMV promoter for oligonucleotide synthesis with common adaptor elements [58] (**Supplementary Table 1**). These regulatory elements were demonstrated to have high levels of transcription in various cell lines [59] and tissues including neural tissues [60,61]. We paired each element with a unique 16bp barcode, individually cloned it into pAAV-Hsp68-nls/mCherry-MPRAe (pAAV-MPRAe), and pooled together in equal molarity (pAAV-MPRAp) to be packaged into AAV to screen viral transduction in target tissues. Since the elements have different sizes, we can detect proper AAV transduction or transcription using polymerase chain reaction (PCR) of genomic DNA or complementary DNA (cDNA), followed by visualization of 3 DNA bands with electrophoresis. Alternatively, these regulatory elements can drive transcription of a nuclear fluorophore which we visualize with immunofluorescence **(Fig 1B)**.

Positive controls

We selected 10 positive controls from Nguyen *et al*. [18] and 20 positive controls from Kheradpour *et al.* [17] to maximize the RNA:DNA ratio, making the enhancer most likely to regulate expression within our assay. For the Nguyen *et al.* enhancers, the simulated neural cell samples were ignored in favor of those enhancers with a higher baseline expression rate. In all cases the positive control enhancers were identified in the cell type similar, but not identical to, the cell type or tissue of interest. We selected candidate brain enhancers from cultured mouse cortical neurons, candidate liver enhancers from HEPG2 cells, and candidate HMC3 enhancers from K562 cells.

Negative controls
We chose 10 negative control enhancers to be the sequences that displayed very low RNA:DNA ratios in cultured mouse cortical neurons from Nguyen *et al.* [18] In addition, we generated a set of 30 random sequence enhancers to create candidate negative control enhancers that were the same length as positive control enhancers with varying GC content (10 each of 30%, 50%, and 70% GC).

Evaluating necessity of MEF2C binding for enhancer activity:
MEF2C is a transcription factor that has been shown to play multiple roles in the cortex and striatum, including driving interneuron morphological maturation [62] and regulating cortical excitatory-inhibitory synapses [38,39], and its up-regulation has been implicated in schizophrenia [63] and reduced vocalization abilities due to its role in repressing dendritic spine development in striatal neurons [39]. These many important roles of Mef2c in the cortex and striatum suggest that the binding of this transcription factor may be necessary for enhancer activity, but this hypothesis was previously infeasible to test because cell lines cannot fully capture cortex and striatum transcriptional regulatory programs. We therefore designed sequences for our WhAMPRA to directly evaluate the necessity of the MEF2C motif for enhancer activity in the brain.

We identified candidate brain-specific enhancers with candidate MEF2C binding sites and designed sequences that compared their activity to the activity of the same sequences without the candidate MEF2C binding sites. To identify candidate brain-specific enhancers, we used the ATAC-Seq / DNase-Seq Pipeline [64] with mm10 [65] and default parameters except for an irreproducible discovery rate (IDR) threshold of 0.1 [66] to call peaks in each mouse motor cortex and striatum replicate from GSE161374 [67]. We used liver as our outgroup tissue for defining brain-specific candidate enhancers, so we repeated this process for the postnatal liver ATAC-seq data from GSE172770 [68–70]. For each of cortex and striatum, we concatenated the peaks from each replicate with the peaks from the liver replicates, merged all peaks within fifty base pairs of each other using mergeBed from bedtools version 2.26.0 with option -d 50 [71], removed parts of peaks overlapping protein-coding transcripts from mouse GENCODE version 15 [72] using subtractBed from bedtools version 2.26.0 with default parameters [71], added 200bp to the end of each peak using slopBed from bedtools version 2.26.0 with option -b 200 [71], and obtained the numbers of reads in each of these regions. Next, for each of the motor cortex and striatum, we obtained peaks that were significantly stronger in each brain region relative to the liver using DESeq2 with default parameters [73], where our cutoffs were fold-change > 1 and adjusted p-value < 0.05. We then obtained the peaks that are significantly stronger in the motor cortex relative to the liver that overlapped peaks significantly stronger in the striatum relative to the liver using intersectBed from bedtools version 2.26.0 with setting -wa [71]. After that, we obtained the sequences of the subset of these peaks that are likely to be enhancers by running closestBed from bedtools version 2.26.0 with options -d -t first and the protein-coding transcripts from mouse GENCODE version 15 [72], removing peaks whose distance from the closest transcript was less than 20kb away, and obtaining the sequences of the remaining peaks with fastaFromBed from bedtools version 2.26.0 [71]. Finally, we obtained the dinucleotide content of these sequences using fasta-get-markov from the MEME Suite with -m 1 [74].

To identify candidate MEF2C binding sites in the brain, we first downloaded MEF2C cortical neuron ChIP-seq data from GSM1629386 and the corresponding input data from GSM162938 [46] and processed the data using the AQUAS Transcription Factor and Histone ChIP-Seq processing pipeline [75] with mapping reads to mm10 [65] and default parameters. We found that the data was high-quality (NSC = 1.09, RSC = 1.20) [68,76], and we obtained 19,418 IDR [66] reproducible peaks across pseudo-replicates (We divided the mapped reads into two pseudo-replicates and used reproducible peaks across them because there was only one biological replicate.). We next identified MEF2C motifs within our brain-specific candidate enhancers by running FIMO [77] on their sequences with the background set to their dinucleotide content and the motif set to the MEF2C motif with ID M4467_1.02 from CIS-BP [78,79]. We then identified the MEF2C peaks overlapping the motif hits, which were our candidate MEF2C binding sites, within our brain-specific candidate enhancers using intersectBed from bedtools version 2.26.0 with options -wa and -wb [71].

We prioritized candidate MEF2C binding sites overlapping brain-specific candidate enhancers by finding those whose brain-specific enhancer activity is especially likely or unlikely to be conserved in other mammals. To evaluate this, we first obtained H3K27ac ChIP-seq regions from human and macaque cortex (occipital pole, precentral gyrus, and prefrontal cortex), striatum (caudate nucleus and putamen) [80], and liver [81] by processing the data using the AQUAS Transcription Factor and Histone ChIP-Seq processing pipeline with option -type histone [75]. We found that all datasets were high-quality, with NSC > 1.05 and RSC > 0.8 for every replicate [68,76]. Next, for human and macaque separately, we merged and extended the peaks from different tissues and different biological replicates in the same way that we did for open chromatin. We then identified the number of reads in each species, tissue, biological replicate combination in each histone modification region using featureCounts with default parameters [82] and after that, in each of human and macaque, used these read counts to identify regions that have significantly higher H3K27ac ChIP-seq in brain relative to liver with DESeq2 with default parameters and alpha=0.05 [73]. We then mapped the macaque H3K27ac ChIP-seq regions to human using liftover [83] and found the subset of human orthologs of regions that have significantly stronger H3K27ac ChIP-seq in macaque brain than in macaque liver that overlap human H3K27ac ChIP-seq regions that are significantly stronger in brain than in liver as well as the subset of human orthologs of macaque regions that are not any stronger in macaque brain than macaque liver that overlap human H3K27ac ChIP-seq regions that are not any stronger in human brain than in human liver using intersectBed from bedtools version 2.26.0 [71] with options -wa (-a was the human regions and -b was the human orthologs of macaque regions) and -u.

To leverage the differential H3K27ac ChIP-seq data to prioritize candidate MEF2C binding sites, we also obtained the candidate MEF2C binding site peak summits overlapping our brain-specific candidate enhancers and mapped them to the human hg38 [84] assembly using liftOver [85]. We then obtained the subset of these summit human orthologs overlapping human H3K27ac ChIP-seq regions that are significantly stronger in brain than in liver and human orthologs of macaque H3K27ac ChIP-seq regions that are significantly stronger in brain than in liver using intersectBed from bedtools version 2.26.0 [71] with options -wa and -u. We also obtained the subset of these summit human orthologs overlap H3K27ac ChIP-seq regions that are not any stronger in brain than in liver and human orthologs of macaque H3K27ac ChIP-seq regions that are not any stronger in macaque brain than in liver in the same way.

We additionally wanted to include candidate MEF2C binding sites that are especially highly conserved, as conservation has been associated with function [86–88]. To do this, we first mapped the MEF2C peak summits at the candidate MEF2C binding sites in mouse brain-specific open chromatin to the human hg19 assembly [89] using liftOver [85]. We next mapped these MEF2C peak summit orthologs to the zebra finch taeGut2 assembly [90] using liftOver [85], thereby identifying peak summits that can be mapped between mammals and birds. Then, we mapped the zebra finch orthologs back to hg19 using liftOver and mapped the hg19 orthologs to hg38 [89] using liftOver [85]. We combined the different groups of human orthologs and mapped them back to the mouse mm10 assembly [65] using liftOver [85].

We further narrowed down these candidate MEF2C binding sites to select those that are non-exonic, have a confident MEF2C motif match within sixty base pairs of their summits, and are near a gene known to be involved in the brain. Specifically, we used subtractBed from bedtools version 2.26.0 [71] to remove any of these candidate MEF2C binding sites that overlapped a mouse protein-coding exon from the mouse Gencode version 15 annotation [72]. We next obtained the sequences of the remaining candidate MEF2C binding site peak summits plus or minus sixty base pairs, where we chose a total length of 120 base pairs because we were planning to synthesize 120 base pair sequences, using fastaFromBed from bedtools version 2.26.0 [71]. We then ran FIMO [77] on these sequences with the same settings as before. We selected the candidate MEF2C binding sites that had a MEF2C motif hit within sixty base pairs of the summit with p-value < 0.0001, which left us with sixty-nine candidate MEF2C binding sites.

For each of the sixty-nine candidate MEF2C binding sites, we created two shuffled versions of each sequence, where, for the first, we shuffled either the region matching the MEF2C motif hit and, for the second, we shuffled the region matching the MEF2C motif hit plus or minus five base pairs. We first created sequences with the shuffled MEF2C motif hits within sixty base pairs of the summits of the remaining candidate MEF2C binding sites by obtaining the motif hit sequences with fastaFromBed from bedtools version 2.26.0 [71], shuffling them by running fasta-shuffle-letters from the MEME suite with options -copies 500 and -dna [74], and creating a sequence for each remaining candidate MEF2C binding site, shuffle combination in which we replaced the MEF2C motif hit with one of the shuffles. We next identified motif hits within these modified sequences by running FIMO [77] on them with the same settings as before except that we included all motifs from CIS-BP version 1.02 from human, macaque, mouse, and zebra finch. We used the results from FIMO to identify motif hits within the shuffled sequences that overlapped motif hits in the original sequences by running intersectBed from bedtools version 2.26.0 with options -wa and -u from bedtools version 2.26.0 [71]. Then, for each of the sixty-nine candidate MEF2C binding sites, we selected the shuffled sequence with the highest motif hit p-value (worst MEF2C motif match) in the region corresponding to the MEF2C motif hit in the original sequence. (If there had been a shuffled sequence with no motif hits, we would have selected that.) We repeated this process for MEF2C motif hits plus or minus five base pairs.

To create our final set of sequences, from the sixty-nine candidate MEF2C binding sites, we removed those with multiple motif hits with p-value < 0.0001, those whose closest gene protein-coding gene was over 350kb away, those whose most significant motif hit p-value in the selected shuffled MEF2C motif hit was lower than the most significant MEF2C motif hit p-value in the original sequence, and a subset of the candidate MEF2C binding sites that were initially selected only because their summits mapped from mouse to zebra finch. To select the subset of candidate MEF2C binding sites that were initially selected only because their summits mapped to zebra finch, we first removed those that were over 100kb from the nearest gene. We determined distances between candidate MEF2C binding sites and genes using closestBed from bedtools version 2.26.0 with option -d [71] and the protein-coding transcripts from mouse GENCODE version 15 [72]. We then kept those that overlapped candidate enhancers that decrease in activity in response to DNA damage, where we mapped those candidate enhancers from mm9 to mm10 using liftOver [85] and identified our MEF2C binding sites that overlapped them using intersectBed from bedtools version 2.26.0 with options -wa and -u [71]. We then kept an additional random fourteen out of the eighteen remaining candidate MEF2C binding sites that were initially selected only because their summits mapped to zebra finch. Finally, we removed all selected candidate MEF2C binding sites that overlapped IDR reproducible motor cortex peaks from seven-week-old mice that were over 1kb long, as we thought these would be unlikely to have enhancer activity without their surrounding sequence, and we identified such overlapping peaks using intersectBed from bedtools with options -wa and -wb [71].

We obtained the orthologs of our final set of twenty eight candidate MEF2C binding sites in human, macaque, zebra finch, and Egyptian fruit bat. We obtained the human and macaque orthologs by mapping the final set of candidate MEF2C binding sites to the human hg38 [89] and macaque rheMac8 [91] assemblies, respectively, using liftOver [85]. We obtained the zebra finch orthologs by mapping the human orthologs to the human hg19 assembly [84] with liftOver [85] and then mapping the outputs from that to the zebra finch taeGut2 assembly [90] with liftOver. We obtained the sequences of the mouse, human, macaque, and zebra finch orthologs using fastaFromBed from bedtools version 2.26.0 [71]. We obtained the Egyptian fruit bat orthologs by finding the Egyptian fruit bat sequences in the Raegyp2.0 assembly [92] that corresponded to the mouse sequences with BLAT [93], as there were no liftOver chains mapping mammalian or avian assemblies to an Egyptian fruit bat assembly. We included the sequences of all of these orthologs as well as the corresponding shuffled sequences in the massively parallel reporter assay (MPRA).

<u>Cortical and Striatal Enhancers near Genes Associated with Vocal Learning</u>
Vocal learning is a complex trait that has evolved independently in multiple clades of birds and mammals [94], serving as a useful trait for the study of the genetic mechanisms underlying the evolution of fine-motor behavior and the connection between genotype and phenotype as a whole. As part of this array, we leveraged epigenomic data for candidate regulatory enhancers

that are both broadly conserved across mammals and proximal to genes known to be associated with vocal learning and disorders of human speech.

First, we generated a set of 57,179 candidate cortical enhancers through intersecting (1) frontal cortex and middle frontal gyrus DNase peaks (ENCODE IDs ENCSR000EIY, ENCSR000EIK, and ENCSR318PRQ) [69], which were from re-processing the data using the ATAC-Seq / DNase-Seq Pipeline [64] with default settings except for -enable_idr -dnase_seq; (2) motor cortex H3K27ac ChIP-seq regions [80], which were from re-processing the data as described above; and (3) a series of brain and liver candidate regulatory enhancers found to be conserved across a wide array of mammalian species [6,80,81]. Second, we generated a set of 53,401 candidate striatal enhancers through intersecting (1) DNase peaks derived from human striatum (ENCODE IDs ENCSR015BGH and ENCSR493VDS [69], which were from re-processing the data as described in our previous work [14]; (2) H3K27ac ChIP-seq regions derived from human caudate and putamen [80], which were from re-processing the data as described above; and (3) a series of brain and liver candidate regulatory enhancers found to be conserved across a wide array of mammalian species [6,80,81]. We found that 39,307 of the 71,273 total candidate regulatory enhancers from both sets (55%) overlapped H3K27ac ChIP-seq regions and DNase peaks in both cortex and striatum and were conserved across multiple mammalian species.

Out of those, we chose a subset of candidate enhancers proximal to genes previously shown to be associated with human apraxia of speech, dyslexia, and stuttering [95–112]. In order to select for likely regulatory enhancers while excluding promoters, we included candidate regulatory enhancers within 200,000 bp of transcription start sites (TSSs) of genes in this set while excluding those within 5,000 bp of TSSs of this gene set. This resulted in a set of 23 candidate enhancers with orthologs in up to 7 species (human (hg38), chimpanzee (panTro4), rhesus macaque (rheMac8), inbred mouse (mm10), Egyptian fruit bat (Raegyp2), little brown bat (myoLuc2), and Zebra finch (taeGut2))—which included 3 confirmed vocal learners; human [113], Egyptian fruit bat [114,115], and Zebra finch [116]—summing to a total of 144 vocal learning-associated enhancer sequences. Sequence orthologs identified across species via BLAT [93] were aligned using PRANK [117] to select the 120 bp segment with the maximal cross-species alignment.

<u>Evaluating effects of AD-associated variants</u>
We downloaded AD GWAS summary statistics from Lambert *et al.* [41] and visualized the GWAS p-values alongside brain cell type-specific H3K27ac ChIP-seq signal tracks and peak calls from our previous work in the Integrated Genomics Viewer (IGV) [118,119]. We selected SNPs to include on the array based on multiple criteria. We selected 15 SNPs that met one or more of the following criteria: (1) the SNP had a significant multiple hypothesis corrected GWAS p-value, (2) the SNP overlapped with H3K27ac peaks in neurons and microglia and, if so, whether it is present in a H3K27ac signal dip [120], (3) the SNP is a sentinel SNP in the AD associated haplotype block or is in high linkage disequilibrium with a sentinel SNP, (4) the SNP disrupts motifs for transcription factors which are important for neuronal and microglial function or highly expressed in neuron and microglia such as SPI1 (Pu.1), EGR1, MEF2, FOXA1, FOXA2 [121], and (5) the SNP had an eQTL association with expression of well-studied AD associated genes that have been shown to be highly expressed in microglia such as BIN1 [122] and SPI1 [123]. In addition,

we selected 12 SNPs that overlapped a human ortholog of a differential H3K27ac peak identified in the brain of the CK-p25 mouse model of AD [123]. We identified human orthologs of the mouse differential peaks using liftover with default settings [85]. For each selected SNP, we included two MPRA enhancer sequences in the array, one carrying the reference allele and the other carrying the alternative allele. In addition, for some cases where the SNP disrupted transcription factor binding site (TFBS) motifs for AD-associated TFs (SPI1, EGR1, MEF2, FOXA1, and FOXA2), we also included a third enhancer sequence that carried a randomly shuffled version of the motif. We centered the sequences on the location of the SNP for all included candidates apart from 10 candidates, where we centered the sequence elsewhere such that the full TFBS motif could be included for the TFBS disruption sequence variants.

### Experimental Design

Library Cloning

We synthesized a new MPRA plasmid construct, pAAV-Hsp68-nls/mCherry-MPRAe (pAAV-MPRAe), based on the previous STARR-seq expression vectors [18] using VectorBuilder Cloning Services. The vector contains AAV2 inverted terminal repeats (ITRs) for use with the PHP.eB packaging system [124] or other AAV2 serotypes, an Hsp68 minimal promoter (with cloning sites at each end to allow for changing of the minimal promoter if necessary), minimal promoter DNA barcode site to allow for the combination of minimal promoters within one library, synthetic intron, mCherry with two SV40 Large T antigen nuclear localization signals (PKKKRKVED) on the N- and C- terminals, and the Chloramphenicol resistance gene surrounded by restriction enzyme sites for cloning in the MPRA insert libraries. The Nextera Transposase binding sequences (Illumina) are located in the plasmid on either end of cloning sites to enable amplification of future barcode libraries (**Supplementary Fig. 1**).

We synthesized the three cross-tissue (CT) positive controls (**Supplementary Table 1**) with strong viral regulatory elements for the MPRAp library in gBlocks by IDT. The insert library (MPRAi; **Supplementary Table 2**) sequence fragments were synthesized by Agilent Technologies. MPRAi and the cross-tissue positive control sequences followed the MPRA Insert Library Template (**Supplementary Fig. 1**) of 28 bases of common sequence for 5’ cloning, candidate enhancer sequence, 27 bases of common sequence linker, 27 bases of DNA barcode, and 35 bases of common sequence for 3’ cloning. We amplified 5 pmol of MPRAi library with Herculase II Fusion Polymerase (Agilent Technologies, #600675), purified it with AMPure XP (Beckman Coulter, #A63881) beads at a ratio of 1.8x, and eluted it in Elution Buffer (EB, Qiagen #19086).

We digested the amplified MPRAi insert library, the CT positive control gBlocks, and the pAAV-MPRAe vector at restriction sites SgsI and SfaAI (**Fig.1A**; **Supplementary Fig. 1**; Thermofisher, #FD1894, #FD2094) and then purified them using QIAEX II kit (Qiagen, # 28704). We dephosphorylated the 5’ ends of linearized pAAV-MPRAe with Shrimp Alkaline Phosphatase (Affymetrix, #78390) to prevent religation of the linearized vector. We ligated the MPRAi library and each CT positive control insert with T4 DNA ligase (NEB, #M0202S) into the linearized pAAV-MPRAe vector and purified them by isopropanol DNA precipitation. We transformed the ligation reactions into MegaX DH10B electrocompetent E. coli cells (Thermo Fisher Scientific,

#C640003). We plated each CT positive control transformation reaction, selected clones, and verified the plasmids with sanger sequencing (Eurofins). We then combined each CT positive control plasmid at equal concentrations for the pAAV-Hsp68-nls/mCherry-MPRAp (pAAV-MPRAp) plasmid library. For pAAV-Hsp68-nls/mCherry-MPRAi (pAAV-MPRAi) plasmid library, a small portion of the transformation reaction was spread on LB agar plates and grown under ampicillin selection. We counted the number of colonies that grew on these plates to verify the transformation efficiency and used the remaining transformation reaction to inoculate LB broth with ampicillin selection to harvest plasmid DNA with an EndoFree Plasmid DNA prep kit (Qiagen, #12362, 12381, or 12391). To check the proportions of the enhancer and barcode libraries, we amplified the pAAV-MPRAi plasmid library with Nextera indexing primers and sequenced it on the Miseq V2 300 cycle kit with paired-end 151 base pair reads and 8 base pair indexing reads (Illumina, MS-102-2002).

Cell Lines

For MPRA library analysis, we purchased human embryonic microglial (HMC3) cells from the American Type Culture Collection (ATCC, cat #CRL-3304) and cultured them according to the manufacturer's guidelines. We purchased AAVPro(R) human embryonic kidney 293 (293T) cells Clontech (cat #632273) for AAV production and cultured them according to the manufacturer's guidelines.

HMC3 MPRAi Plasmid Transfection

We seeded HMC3 cells in a 100 mm dish to be approximately 80% confluent on the day of transfection. We transfected cells with 20 µg of pAAV-MPRAi plasmid library DNA using FuGene 6 transfection reagent at a FuGene6:DNA ratio of 3:1. We incubated cells at 37˚C for 72 hours. We harvested the cells from each dish 72 hours post-transfection and centrifuged them at 500 x g for 5 minutes. We extracted genomic DNA and total RNA using DNeasy Blood & Tissue Kit (Qiagen, Cat #69504) and RNeasy Mini Kit (Qiagen, cat#74104), respectively. We then stored the DNA at -20˚C and the RNA at -80˚C until we were ready to prepare the DNA barcode libraries.

Adeno-Associated Virus Packaging and Titration

We began adeno-associated virus (AAV) production by co-transfecting the pAAV-MPRAp or pAAV-MPRAi plasmid libraries with packaging vectors tTA- iCAP-PHP.eb (pUCmini-iCAP-PHP.eB) and pHelper into AAVpro HEK 293T cells (Clontech, #632273) via Linear 25kDa Polyethylenimine (PEI; Polysciences, #23966-2). We received the packaging vectors, pUCmini-iCAP-PHP.eB, and pHelper, as a gift from Vivianna Gradinaru. (http://addgene.org/; 103005; RRID: Addgene_103005) (Chan et al., 2017). At 72 hours post-transfection, we collected the media supernatant from the cells, stored at 4˚C, and added fresh media. At 120 hours post-transfection, we collected the media supernatant from the cells, combined it with the 72 hour supernatant, and precipitated the AAV particles harvested with 4% polyethylene glycol (PEG 8000, Sigma #P2139) in 2.5M sodium chloride (Thermofisher, #AAJ2161836) overnight at 4˚C. We harvested the AAV particles from the transfected 293T cells 120 hours post-transfection and centrifuged them at 4,000 x g. We lysed the cells by resuspending the cell pellet in SAN Buffer (40 mM Tris, 500 mM NaCl, 2 mM MgCl2 pH 8.0) with

Salt Activated Nuclease at 20 U/mL (SAN enzyme; Arcticzymes, #70900-201) and incubated at 37˚C for 1 hour, with intermittent vortexing. Subsequently, we froze the lysed cells at -80˚C. At 144 hours post-transfection we centrifuged the PEG precipitated media supernatant at 4,000 x g to harvest AAV particles and lysed the pellets by resuspending in SAN Buffer with SAN enzyme at 37˚C for 30 minutes with intermittent vortexing. Afterward, we stored the lysates at -80˚C overnight. We centrifuged the cell lysates and PEG lysates were centrifuged at 2,000 x g and then collected the AAV through an iodixanol gradient via ultracentrifugation (15%, 25%, 40%, 60%; Optiprep, Sigma, #D1556-250ML [125]. We withdrew the 40-60% interface and diluted it with sterile phosphate buffer saline + pluronic F-68 (Thermofisher, #24040032). We concentrated and purified the viruses using Amicon Ultra 15 kDa filter (Fisher Scientific, #UFC910096), eluted them in sterile phosphate buffer saline (PBS), and filtered them for injection through a 0.22 µm syringe filter. We titered the virus using the AAVpro Titration Kit (Clontech, #6233), aliquoted it into LoBind tubes (Eppendorf, cat #0030108434), and stored it at −80°C until injection.

Animals:

All procedures were approved by Carnegie Mellon University Institutional Animal Care and Use Committee (IACUC). We performed molecular and imaging experiments on 3 to 6 month old C57BL/6J mice from The Jackson Laboratory (strain #000664). We used female and male mice. We injected the pAAV-MPRAi library into 8 mice, 4 males and 4 females, for RNA sequencing and 4 mice, 1 male and 3 females, for immunofluorescence experiments (**Fig. 1**). We injected the pAAV-MPRAp library into 2 female mice for immunofluorescence imaging experiments (**Supplementary Table 3**).

We anesthetized the mice using isoflurane at 1-4% until breathing slowed and the animal had no pedal reflex. We injected $7.73 \times 10^{11}$ to $2.28 \times 10^{12}$ vector genomes (vg) total into the retro-orbital cavity. Following the injections, the mice received 0.5% proparacaine hydrochloride ophthalmic solution for comfort and we monitored the mice for any abnormalities or signs of distress after the procedure. We left the virus to incubate for 3-6 weeks before collecting tissue for downstream experiments.

In RNA-seq experiments, we deeply anesthetized the animal with isoflurane until a lack of pedal withdrawal and decapitated it. Immediately following death, we harvested fresh tissues. We sectioned brain tissue with a Leica VT 1200 vibrating microtome at a thickness of 300 µm that was staged in cold, oxygenated artificial cerebrospinal fluid (aCSF). We dissected the primary motor cortex (M1), prefrontal cortex, other frontal cortex (referred to as Cortex throughout this paper), striatum, hippocampus, and hypothalamus from the brain sections, divided each brain region into 2 tubes, flash froze them, and stored them at -80˚C until processing. We also harvested liver, testes, ovaries, and heart immediately following decapitation, minced them into small pieces with a clean razor blade, divided them into 2 tubes, flash froze them, and stored them at -80˚C. Genomic DNA and total RNA were extracted using DNeasy Blood & Tissue Kit (Qiagen, Cat #69504) and RNeasy Mini Kit (Qiagen, cat#74104), respectively. We then stored the DNA at -20˚C and the total RNA at -80˚C until the DNA barcode libraries were prepared for sequencing.

For imaging experiments, we deeply anesthetized mice with isoflurane and confirmed with a negative toe-pinch response, injected with intraperitoneal urethane (50 mg/mL, Acros Organics, cat #A0378229), and performed cardio-thoracic perfusion with 1x PBS followed by 4% paraformaldehyde (approximately 10 mL), after which we harvested tissues and incubated them in 4% paraformaldehyde for 4-12 hours at 4˚C. Following incubation, we washed tissues with 1x PBS to remove PFA and stored them in 1x PBS at 4˚C until ready to process for immunofluorescence staining and imaging.

Immunofluorescence staining and imaging:
We sectioned tissues on a Leica VT1000S vibratome at 80 µm slices and probed them for WhAMPRA expression of the nuclear mCherry reporter, using a standard immunohistochemistry protocol. We stained brain tissues with primary anti-NeuN (Cell Signaling #94403, 1:500) with secondary AlexaFluor 488 (Invitrogen #A11029, 1:500) and primary anti-mCherry (Cell Signaling #43590, 1:500) with secondary AlexaFluor 594 (Cell Signaling #8889, 1:500; Fig. 1B&C). We stained liver tissue with 4′,6-diamidino-2-phenylindole (DAPI, Thermofisher #D1306) and primary anti-mCherry (Cell Signaling #43590, 1:500) with secondary AlexaFluor 594 (Cell Signaling #8889, 1:500). We mounted slices on glass slides (Color Plus Microscope Slides, Cat. #12-550-18) and coverslipped the tissue with ProLong Diamond Antifade Mountant (Thermo Fisher Scientific, cat #P36961). We imaged slides with a Zeiss 880 Laser Scanning Microscope using a spectral analysis camera. We performed all image processing using Zeiss Zen Black software and ImageJ to analyze images.

MPRAi DNA Barcode Library Preparation from Tissue and HMC3 Transfected Cells:
To prepare the DNA barcode libraries for sequencing from extracted total RNA, we treated the RNA with Turbo DNase (Thermofisher Scientific, cat#AM2238) and SUPERase-In RNase Inhibitor (Thermofisher Scientific, cat#AM2694) following the manufacturer's instructions to remove any possible contaminating AAV vector genome DNA. We purified the RNA with the RNeasy MinElute CleanUp Kit (Qiagen, Cat #74104) and quantified concentrations using a Qubit fluorometer. We performed a specific Reverse Transcription (RT) on up to 2 µg of RNA using the SuperScript IV enzyme (Thermofisher, Cat #18090200) and a specific RT primer (GTACAAGAAAGCTGAACGAGAAACG) complementary to the 3' tail of the MPRA transcript before the SV40 polyadenylation signal, followed by an RNA denaturing treatment using 1M NaOH, pH > 10 at 98˚C for 20 minutes. We purified the cDNA for each sample by isopropanol precipitation with GlycoBlue Co-precipitant (Thermofisher Scientific, Cat #AM9515) to help visualize the DNA pellet.

We amplified both genomic DNA and cDNA with dual-indexing primers synthesized from Eurofins following the Nextera tagmentation format as previously published in Preissl et al. Supplementary Table 5 [126] and Phusion High-Fidelity PCR Master Mix (Thermo Fisher Scientific, #F531S). We then purified all samples and concentrated them with the MinElute PCR Purification Kit (Qiagen, cat #28004). We measured the quantity and quality of each sample using the Qubit and Agilent TapeStation.

Sequencing:

To obtain the initial library quality estimates and balance sample representation for deep sequencing, we pooled each sample and sequenced with 5% PhiX (Illumina, #FC-110-3001) on the Illumina MiSeq system using a 150-cycle V3 Kit (Illumina, Cat #MS-102-3001). We rebalanced the library pool prior to deeper sequencing and sent the libraries to be sequenced to (#of Reads/sample) with 30% PhiX on 2 Novaseq S4 flowcells (GenWiz by Azenta). Due to the number of total samples in the project, we used two separate NovaSeq experiments to reach target sequencing depth across samples. To eliminate possible batch effects, we included several high-quality and low-quality samples to be repeated in each sample pool. For all sequencing runs and intermediate steps, we followed the Illumina and GenWiz guidelines.

### Computational Analyses:

Computational Library Processing;

Across each of the sequenced samples (**Supplementary Table 3**), we used a custom program to count the barcode reads at the DNA and RNA levels (arrayProc.2.1.1.py). We searched for close matches to the designed restriction enzyme site and then searched the neighboring bases for matches to the designed barcode. For each sample, we calculated the number of reads with the barcode at the DNA (**Supplementary Table 4**) and RNA (**Supplementary Table 5**) level. We also calculated the proportion of those reads that include a restriction enzyme site and recognizable barcode.

We read, processed, and formatted the library information, sample information, and barcode counts per sample. Over 90% of enhancers were associated with at least 5 barcodes in the final library. For each sample at both the DNA and RNA level, we calculate a number of quality control metrics in addition to the proportion of barcodes detected. We suspect that, due to the low amount of RNA in certain samples, that the dropout could be high. To detect potential dropouts, we compared the RNA-DNA ratios of enhancers in each sample. In some samples, the RNA levels were very low despite strong DNA counts (**Supplementary Fig. 3A**). In other samples, very few enhancers showed lower than expected RNA counts (**Supplementary Fig. 3B**). We summarized the potential dropout tendency with an "outlier score," calculated as negative on tenth the number of log2(candidate enhancer RNA counts/candidate enhancer DNA counts) < 0.3, which tends to be lower if the sample quality is lower (**Supplementary Table 3**). On the basis of this score, we removed all samples from tissues with low amounts of RNA; we think that the removed tissues (hypothalamus, muscle, kidney, lung, heart, ovaries, testes) had low RNA amount because very few enhancers were designed for them. For this reason, we also removed the partial frontal cortex (PFC) samples, which were a test of the assay with low amounts of brain tissue. We additionally eliminated several individual samples based on this score, including the striatum for animal 5 and the hippocampus from animal 6. Finally, we removed the RNA from HMC3 cell sample C, which showed low experimentally measured RNA quality and then low correlation with RNA measured in other HMC3 samples. At the DNA level, we removed the striatum sample for animal 2, which had substantially fewer reads (million reads=2.1 << median 21.6). We also removed HMC3 sample E due to poor DNA correlation with all other samples R=0.81 << median R=0.95)

For the remaining samples, we merged the highly correlated read counts across the two NOVA-Seq runs and the counts across barcodes to create matrices of read counts per sample, enhancer combination. We then used MPRAnalyze [45] to estimate the raw transcriptional activity (alpha) and the normalized transcriptional activity (MAD Score) of each enhancer. We did this analysis at multiple levels: per sample, per tissue, and per tissue type (brain, liver, HMC3). We also calculated the significance of the enhancer activity relative to negative control levels (**Fig. 2A**).

Comparison to Machine Learning Model Predictions:
We used our brain and liver open chromatin prediction models trained using open chromatin data from multiple species (models 8-9) from our previous work [14] to predict brain and liver open chromatin for our WhAMPRA candidate enhancers. Since our models required 500 base pair sequences and our WhAMPRA enhancers were only 120 base pairs, we used biopython version 1.74 [127] to pad each sequence with 190 base pairs of Ns on each side, creating a 500 base pair sequence with the WhAMPRA candidate enhancer in the center. We put the sequences through the models using keras version 1.2.2 [128]. This gave us a prediction for each WhAMPRA candidate enhancer sequence as well as its reverse complement. When comparing these predictions to enhancer activity measured by the WhAMPRA, we took the average prediction made by the forward and reverse complement.

Disruptions of Transcription Factor Binding Sites and SNPs:
We used the MPRAnalyze-calculated MAD score to compare the sequences designed at MEF2C transcription factor binding sites to versions of those sequences where the MEF2 transcription factor binding site or the site along with the surrounding region was shuffled. We used a paired t-test across all samples in the primary motor cortex, other cortex, and striatum to identify cases where disrupting the MEF2 transcription factor binding site motif disrupted enhancer activity (**Supplementary Table 7**).

To compare the enhancer activities of the different alleles and disruptions of candidate AD-associated mutations, we also used a paired t-test of the MAD scores (**Supplementary Table 8**). Here, we used the HMC3 cells and the neural tissues implicated in AD predisposition and progress, but not liver tissue, for the comparison.

**Github**: https://github.com/pfenninglab/WhAMPRA
Token available upon request

### Supplemental Figures:

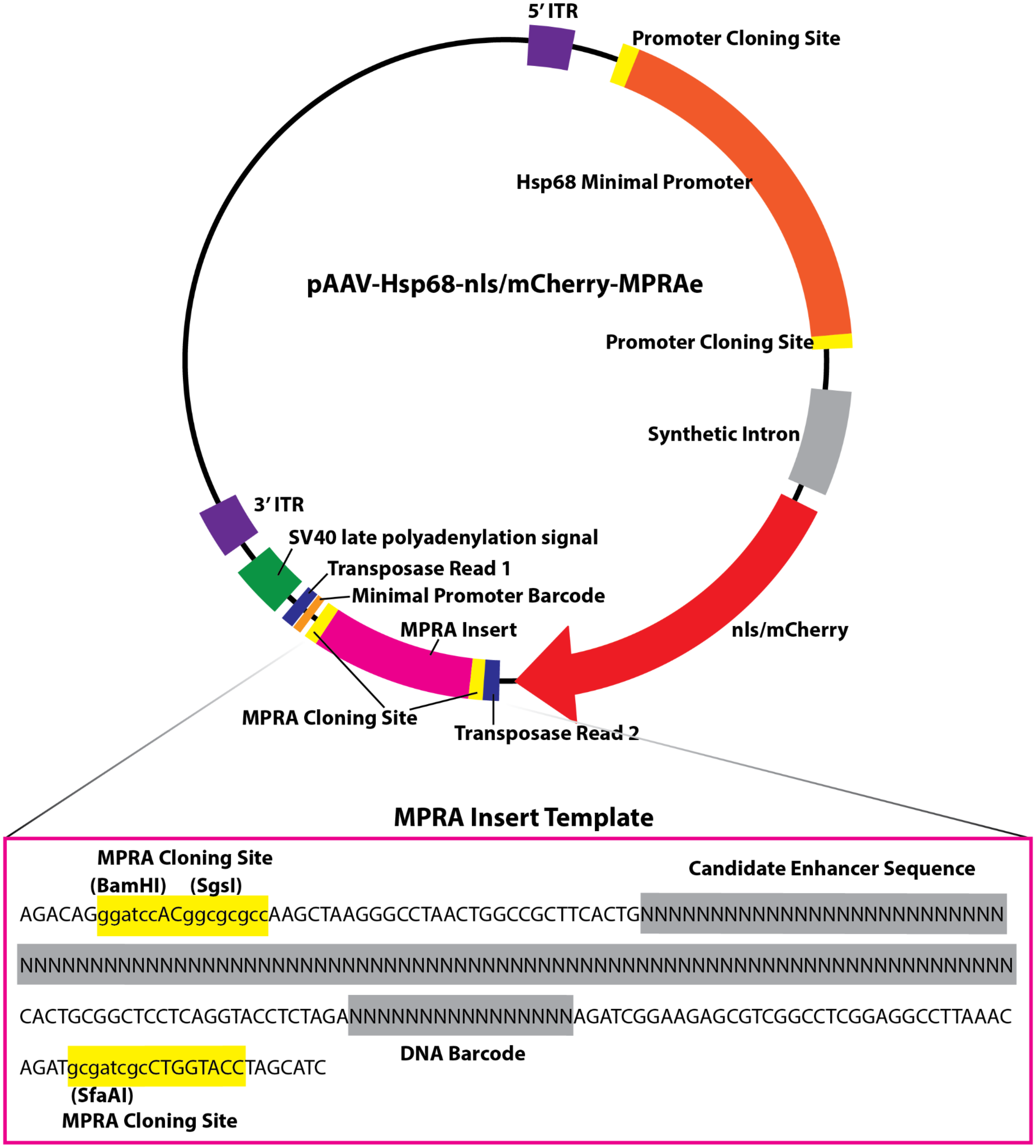


**Supplementary Fig. 1** - MPRA plasmid design

The top of the figure shows a map of pAAV-Hsp68-nls/mCherry-MPRAe (pAAV-MPRAe) design features. The plasmid features listed from 5' to 3': 5' inverted terminal repeat sequences (ITR) for PHP.eB and AAV 2 serotypes, Hsp68 minimal promoter with cloning sites at either end for the flexibility for changing the promoter, synthetic intron, mCherry with two nuclear localization sequences (nls) fused to the protein, Illumina Transposase Read 2 sequence, MPRA insert library with cloning sites at either end, minimal promoter DNA barcode, Illumina Transposase

Read 1 sequence, SV40 late polyadenylation signal (polyA tail), and 3' ITR. The bottom of the figure shows a more detailed view of the MPRA insert template used to design the libraries. The sequences listed are the same for all oligos, while the gray shaded areas vary with the different enhancers and DNA barcode sequences. The yellow highlighted areas show the cloning sites, with the restriction enzyme digestion sequences in lower case letters.

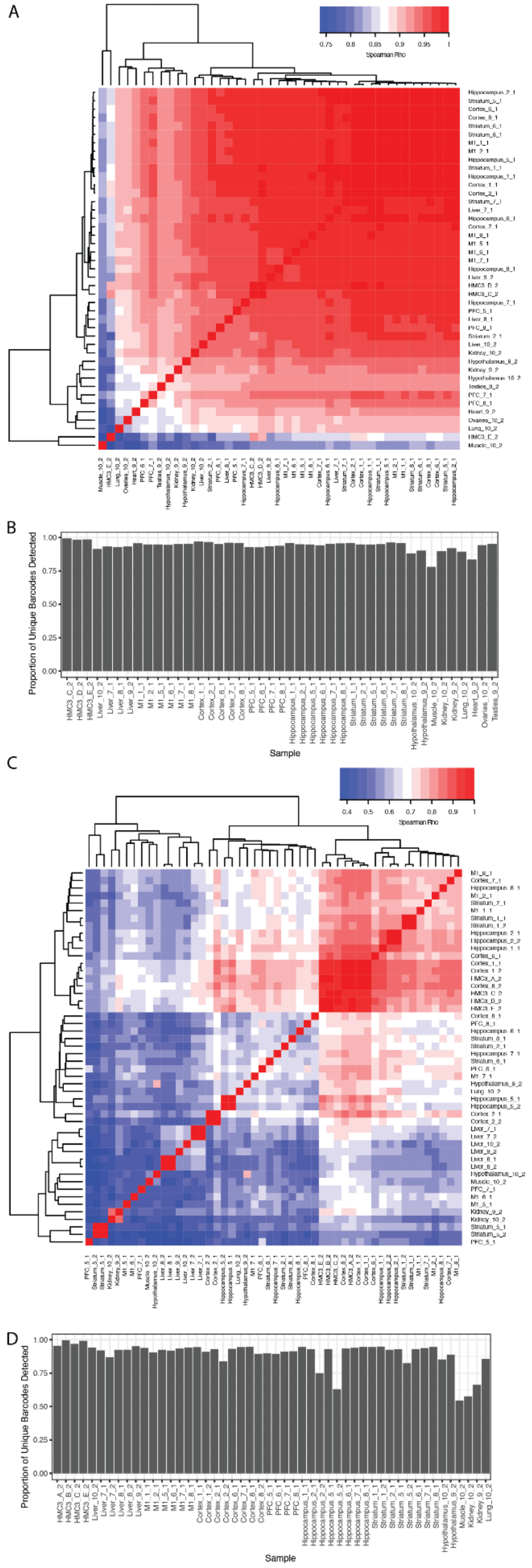
A
Spearman Rho
B
Proportion of Unique Barcodes Detected
Sample
C
Spearman Rho
D
Proportion of Unique Barcodes Detected
Sample

**Supplementary Fig. 2** - Correlation of DNA and RNA barcode measurements across tissues. The Spearman correlation across all samples for the plasmid DNA reads (A) for the RNA reads (C). For each sample, we plot the proportion of barcodes detected in that sample relative to detected in any samples (3,983) for both DNA (B) and for RNA (D).

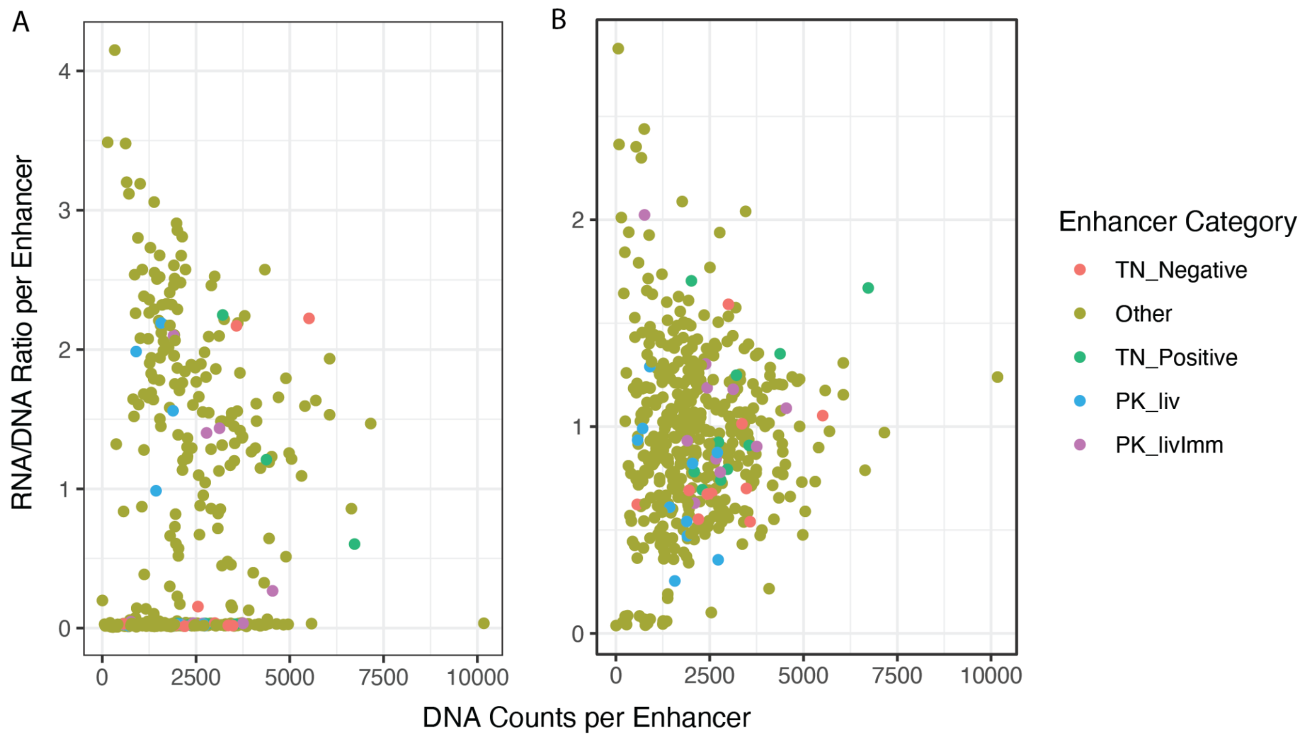


**Supplementary Fig. 3** - Enhancer dropout in low quality samples.
The RNA:DNA ratio is plotted relative to the number of barcode reads from the plasmid DNA (thousands). Two samples, striatum from animal 2 (A) and striatum from animal 5 (B) are displayed. Each enhancer is colored based on its design group. Substantial numbers of positive control and other candidate enhancers appear to have dropped out of the striatum from animal 2.

### Supplementary Tables:

All Supplementary Tables can be found at
http://daphne.compbio.cs.cmu.edu/files/ikaplow/WhAMPRASupplement/

**Supplementary Table 1** - Sequences for the MPRAp Library
The sequences for the cross-tissue positive control library (MPRAp). The columns list the synthesized DNA sequence for cloning, the enhancer sequence, and unique 16bp barcode sequence.

**Supplementary Table 2** - Sequence and Annotation of the MPRAi Library
There is a column for the enhancer sequence, barcode, full oligonucleotide sequence synthesized, source of the sequence, and whether or not the sequence was detected at each of the RNA and DNA levels. The source of sequence refers to the topic of the question it was intended to address (AD - sequence with Alzheimer's disease variant; GC - Randomly generated sequences with specific GC contents; MEF2C - MEF2C-bound, MEF2C-bound with motif shuffle, or ortholog of MEF2C-bound sequence; VL - sequence implicated in vocal learning evolution; PK_liv - sequences with liver specific enhancer activity from [17]; PK_livImm - sequences with enhancer activity in liver and K562 from [17]; TN_cort - sequences with high baseline activity in cultured cortical neurons from [18]; TN_neg - sequences with low baseline activity in cultured cortical neurons from [18].

**Supplementary Table 3** - Sample and Quality Control Information
For each sample, we calculated the number of reads with the barcode at the DNA and RNA levels. We also calculated the proportion of those reads that included a restriction enzyme site and recognizable barcode.

**Supplementary Table 4** - DNA Counts
The number of each barcode measured at the DNA level.

**Supplementary Table 5** - RNA Counts
The number of each barcode measured at the RNA level.

**Supplementary Table 6** - Machine Learning Model Predictions
For each enhancer, we provided the predicted activity or open chromatin level across the different cell types and tissue assayed.

**Supplementary Table 7** - MEF2 Binding Site Disruption
The results of running a paired t-test between each candidate MEF2-binding enhancer and the versions where the MEF2 motif is disrupted by shuffling the nucleotides.

**Supplementary Table 8** - Disruption of Candidate AD-Associated SNPs at Candidate Enhancers
The results of running paired t-tests between each of the reference and alternate alleles of candidate enhancers associated with AD predisposition.
